# Accelerated aging induced by stress in experimental murine ocular hypertension

**DOI:** 10.1101/2022.05.01.490248

**Authors:** Qianlan Xu, Cezary Rydz, Viet Anh Nguyen Huu, Lorena Rocha, Claudia Palomino La Torre, Irene Lee, William Cho, Mary Jabari, John Donello, Robert N. Weinreb, David C. Lyon, Won-Kyu Ju, Andrzej Foik, Dorota Skowronska-Krawczyk

**Affiliations:** Department of Physiology and Biophysics, Center for Translational Vision Research, School of Medicine, University of California, Irvine, Irvine, CA; Viterbi Family Department of Ophthalmology, Hamilton Glaucoma Center and Shiley Eye Institute, School of Medicine, University of California, San Diego, La Jolla, CA; Department of Anatomy and Neurobiology, School of Medicine, University of California, Irvine, Irvine, CA, USA; International Centre for Translational Eye Research, Institute of Physical Chemistry, Polish Academy of Sciences, Warsaw, Poland; Department of Physiology and Biophysics, Department of Ophthalmology, Center for Translational Vision Research, School of Medicine, University of California, Irvine, Irvine, CA

**Author notes:** equally contributed (Qianlan Xu, Cezary Rydz). Corresponding author **Dorota Skowronska-Krawczyk, PhD**, Department of Physiology and Biophysics, Department of Ophthalmology, Center for Translational Vision Research, University of California Irvine, Gavin Herbert Eye Institute, 837 Health Sciences Rd, Irvine, CA 92617.

**Keywords:** stress response, aging, retinal ganglion cells, IOP, senescence

## Abstract

Aging, a universal process that affects all cells in an organism, is a major risk factor for a group of neuropathies called glaucoma, where elevated intraocular pressure is one of the known stresses affecting the tissue. Our understanding of molecular impact of aging on response to stress in retina is very limited, therefore we developed a new mouse model to approach this question experimentally. Here we show that susceptibility to response to stress increases with age and is primed on epigenetic level. We demonstrate that program activated by hypertension is similar to natural aging, and that one of the earliest pathways activated upon stress is senescence. Finally, we show that multiple instances of pressure elevation cause accelerated aging of young retina as measured on transcriptional and epigenetic level. Our work emphasizes the importance of early diagnosis and prevention as well as age-specific management of age-related eye-diseases, including glaucoma.

## INTRODUCTION

Aging is a complex process for which distinct molecular aspects contribute to age-related tissue dysfunction. Systematic analysis of epigenomic and transcriptomic changes across mouse tissues during aging identified several recurring processes including interferon alpha response, IL6-JAK-STAT3 signaling, complement and other components of innate immune response being gradually upregulated with age.^1^ These changes are conserved as the molecular signatures of aging and are consistently upregulated mice and other vertebrates in the absence of pathogens. One aspect of age-related changes is sterile inflammation, commonly termed “Inflammaging”, has been reproducibly observed in many laboratories.^2,3,4,5^

One type of age-related disease is glaucoma, an eye disease in which there is interaction of multiple genetic and environmental factors^6,7^. It is characterized by progressive neurodegeneration of the optic nerve that if untreated leads to morphological changes, functional decline, and eventual blindness. Of all the risk factors that have been reported for glaucoma, age is among the strongest and the most consistent. Other risk factors, including elevated intraocular pressure (IOP), family history, and high myopia exhibit more variability. Because of the rapid increase in aging populations worldwide, current estimates show that the number of people with glaucoma (aged 40-80) will increase to over 110 million in 2040.^8^

Until now, lowering IOP is the only approved and effective treatment paradigm. However, many treated glaucoma patients continue to experience deterioration of vision and progress to blindness, highlighting the critical need to understand its underlying mechanisms. ^6,9,10^

Pathophysiological stress related to elevated IOP induces a broad spectrum of changes in the connective tissues and blood vessels that are central to the function of the retina. They also induce molecular changes in retinal cells, including retinal ganglion cells (RGCs). We have shown previously that with experimental ocular hypertension, RGCs respond to stress by expressing *p16Ink4a*, assume a senescent phenotype and secrete molecules known as senescence associated secretory phenotype (SASP) such as IL6. Similarly, we have observed an increase of senescent cells in our mouse model of ocular hypertension and in human glaucomatous retinas.^11^ Senescence is a type of damage response that develops in cells experiencing irreparable stress for a sustained period of time, and it is commonly observed in all aged tissues, including the eye. ^12,13^ Therefore, we investigated the extent to which aging contributes to IOP-related senescence induction, stress and RGC death upon IOP elevation.

In this work, we describe functional and molecular changes in the aging retinas, showing changes characteristic of inflammaging, as found in other tissues^2,14,15^, and strong downregulation of retinal lipid metabolism. We show that aged retinas are more sensitive to mild IOP elevation and that old RGCs express significant numbers of senescence markers upon IOP-associated stress. Further, we show that senescence is significantly upregulated in old retinas among the first group of activated pathways upon IOP elevation, and that RGC survival can be promoted using a senolytic drug immediately after IOP elevation. Next, we demonstrate that aged retinas are prone to respond to the insult more robustly than young tissues and that this correlates with the pre-opened enhancers. Finally, we show that multiple mild instances of IOP elevation in young animals preprogram the tissue to respond more strongly to stress, similar to the response of the aged tissue. Our DNA methylation analysis revealed that multiple mild stresses on the young retinas accelerated DNA methylation clocks, suggesting the direct influence of repetitive stress on retina aging.

Our results collectively suggest that throughout the life of the animal, retinal cells accumulate the competence on the epigenetic level to respond to stress faster and stronger, eliciting more damage to the tissue in the form of the inflammation and senescence, including the secretion of SASPs and destabilization of the extracellular matrix. These results might explain the difficulties in managing the responses to glaucomatous insults, such as elevated IOP, in aged individuals and suggest novel approaches to manage age-related eye diseases, including glaucoma.

## RESULTS

### Functional and molecular changes in aging retina

First, the impact of aging on visual functions in young (3-month-old) and old – (18months old) mice was measured using several functional assays.

The quantitative optomotor response (OMR) analysis was performed to measure contrast sensitivity in scotopic (nighttime) and photopic (daytime) light level conditions. In this assay, a mouse is placed on a platform where it can move and track a stimulus – rotating pattern displayed on a screen **(Figure 1A, top)**. Response evaluation is automatic and presented as an OMR index (ratio of correct/incorrect). Contrast sensitivity was defined as the inverse of contrast threshold for OMR and measurements were performed using an automated setup (PhenoSys GmbH) **(Figure 1A, Supplementary Figure 1A)**. Our data shows that old animals have significantly lower contrast sensitivity in both scotopic and photopic conditions relative to young animals **(Figure 1A)**. For example, in low luminance of ∼0.03 lux (nighttime light levels) the tracking behavior of aged animals was decreased already at 100% contrast. In daytime light levels 18-month-old animals exhibited significantly decreased tracking response already at 50% contrast when compared to the younger animals **(Supplementary Figure 1A)**.

**Figure 1.**
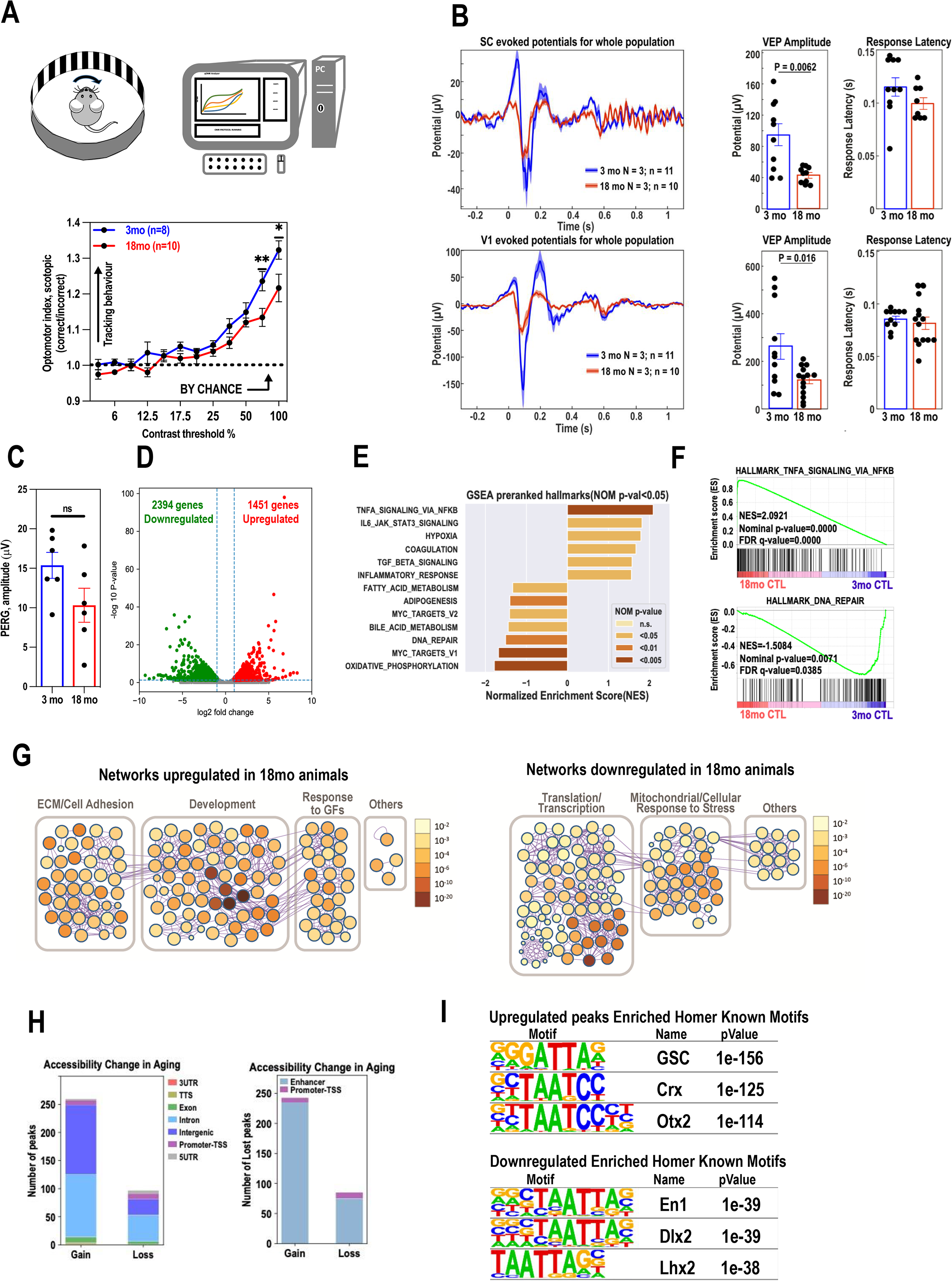
Functional and molecular changes in aging retina. **A**. Schematic of the optomotor response (OMR) instrument. (top) OMR at various pattern contrast (bottom). Evaluation of response is automatic and presented as OMR index (ratio of correct/incorrect). Statistically significant tracking obtained using one tailed test. Mean and S.E.M. are shown. \**P* < 0.05, \*\**P* < 0.01. **B**. Visually evoked potential recordings. VEPs were recorded in Superior Colliculus (SC) left-top, and in primary visual cortex (V1) left-bottom. VEP amplitudes and Response latencies (right-top, right-bottom) are presented. **C**. PERG performed on 3-month-old (n=6 eyes) and 18-month-old (n=6 eyes) wild type mice shows no significant change in amplitude in aged animals. (Unpaired, two-tailed t-test). **D**. Volcano plots of differentially expressed genes between 3-month-old and 18-month-old non-treated wildtype mouse retinae. Each dot represents one gene with -log10(adjusted P-value) versus log2foldchage. Dots in grey represent no significant genes while the dots highlighted in red and green respectively represent up-regulated and down-regulated DEGs. n=2 **E**. The bar plot showed normalized enrichment scores (NES) of all defined significant enriched (nominal p-value <5%) hallmark gene sets from pre-ranked GSEA in 18-month-old non-treated wildtype mouse retinae compared to the 3-month-old retinae. n = 2 biological replicates. NES determines the magnitude of enrichment of each hallmark gene set across all analyzed gene sets. The statistical significance in nominal (NOM) p-value were indicated by a discrete color scale. **F**. GSEA enrichment plots of selected top-ranked hallmark pathways **G**. Networks of enriched pathway from up- and down-regulated DEGs during aging using Metascape. Each node in the figure represents one enriched ontology term. Node size indicates the number of genes in each term. The statistical significance in FDR q-value were shown by a discrete color scale for each node. All the enriched terms were further bounded and annotated as more general classes. **H**. Stacked bar plots showed up- and down-regulated accessible regions with cutoffs at 0.05 in FDR and two-fold change of normalized ATAC-Seq signal. Specific genomic regions were color coded. **I**. Motif analyses using tool Homer identified potential regulators during retina aging. Regions of accessibility gain, or loss were enriched in Homer Known Motifs. The table showed top ranked potential transcription factors in significance by motif analysis of ATAC-Seq data in comparison between 3-month-old and 18-month-old mouse retinae.

Next, we evaluated age-related effects on the integrity of the visual pathway by recording visually evoked potentials (VEP) in young and old animals. The recordings were performed in the superior colliculus (SC) and the primary visual cortex (V1). In both regions there was a significant reduction in VEP in 18-month-old mice (red trace) compared to the 3-month (blue trace) animals. For example, in the SC, the VEP amplitude dropped by 56% from 95 ± 14 µV in 3-month to 42 ± 4 µV in 18-month-old animals (P = 0.006). In V1, the drop was 60%, from 263 ± 54 µV in 3-month to 106 ± 12 µV in 18-month-old animals (P = 0.016). Ion the other hand, response latencies were comparable between age groups in both regions (115 ± 7 ms vs. 99 ± 6 ms in the SC; and 85 ± 3 ms vs.81 ± 6 ms in the V1; **(Figure 1B)**.

To examine the extent to which the observed decrease of visual function can be attributed to the loss of RGC activity, the pattern electroretinography (PERG) responses were measured. PERG signals (OS) from six 3-month and six 18-month-old animals were recorded. Our findings demonstrate no significant difference in PERG amplitudes between young (mean, 15.37 ± 4 μV [SD]; *n* = 6 eyes) and old animals (mean, 10.32 ± 5.29 μV]; *n*=6 eyes) (t=1.87, p=0.09) **(Figure 1C)**.

To investigate whether the molecular changes in the aging retina can explain the observed functional decline of the visual system we performed a whole retina mRNA sequencing on retinas isolated from both 3-month and 18-month-old mice. We found that there were 1451 genes significantly up-regulated (log2 fold≥1, FDR≤0.05) and 2394 genes significantly down-regulated (log2 fold≤-1, FDR≤0.05) upon natural aging **(Figure 1D)**. The pre-ranked Gene Set Enrichment Analysis (GSEA) revealed a dysregulation in several conserved pathways (MsigDB hallmark collection version v7.5.1) in aged animals.^16^ In particular, we found a significant upregulation of inflammatory response (NES=1.56), NF-kB regulated TNF alpha signaling (NES=2.09) and IL-6 Receptor Signaling (NES=1.82) and a significant downregulation of DNA repair (NES=-1.51), fatty acid metabolism (NES=-1.34) and oxidative phosphorylation pathways (NES= −1.78) in the 18-month-old mice relative to the 3-month-old animals **(Figure 1E, F)**.

Metascape enrichment network^17^ visualized the intra- and inter-relationships among the up- and down-regulated gene pathways in natural aging (3-month-old versus 18-month-old) **(Figure 1G, Supplementary Figure 1B)**. capturing the clustered upregulation of processes such as extracellular matrix organization, response to growth factors and general development. Downregulated clusters included stress-related mitochondria organization and translation.

To compare transcriptomic data with epigenetic changes during natural aging, we performed an Assay for Transposase-Accessible Chromatin with high-throughput sequencing (ATAC-Seq) analysis on 3-month-old (N=6) and 18-month-old (N=6) mouse retinas. The epigenetically affected genes were determined by the significant differential chromatin accessibility in both promoter-TSS and the nearest enhancer regions **(Supplementary Figure 1C)**.

The open chromatin regions or significant reads coverage (q-value ≤ 0.05) of ATAC-Seq data were identified by peak calling using MACS2 (Model-based Analysis for ChIP-Seq, version 2.2.7.1). The chromatin accessibility of the young and old retina samples was compared by using differential analysis tool edgeR in the count matrix built from the consensus list of detected open regions from all samples with significance cutoff at 0.05 in FDR q-value. 260 regions gained accessibility in 18-month-old retina while 97 gained accessibility in 3-month-old mouse retina **(Figure 1H)**. Majority of the age-related epigenetically remodeled regions were not on the promoters of the genes. Similarly, loss of the accessible regions only partially (∼10%) overlapped with promoters of the genes. Transcription factor (TF) motif analysis indicated enrichment of unique sets of homeodomain transcription factors in both groups **(Figure 1I, Supplementary Figure 1D)**. In particular, predicted binding sites for Crx and Otx2, TFs involved in development and maintenance of the photoreceptor cells are highly enriched.

### Aged retina is more sensitive to mild intraocular pressure elevation

In our previous works we have demonstrated that intraocular hypertension of 90 mmHg, non-physiological levels of pressure, induced ∼50% loss of RGCs.^11,18^ Therefore, in search of a better system that would mimic the glaucomatous insult, the effects of different levels of IOP elevation on RGC survival were quantified, by unilaterally elevating IOP in 3-month-old C57BL/6J to 30, 50, or 90 mmHg for 1 hour. The contralateral eye served as a healthy sham control. At day 7 post-IOP, retinas were dissected, fixed, and stained using the antibody against Brn3a or RBPMS, specific markers for RGCs. In comparison with control, we found that only IOP elevated to 50 and 90 mmHg pressure induced a significant decrease in the RGC count. In addition, there was a direct correlation between the magnitude of RGC loss with the level of applied IOP. Mild IOP elevation of 30 mmHg had no impact on RGC count in 3-month-old animal after 7 days. To explore whether simple needle puncture can contribute to RGC loss, 7mm Hg and 15mmHg pressure cohorts were tested. At 7 days post-procedure, analysis of the immunostained retinal flat mounts revealed no decrease in RGC count in procedural eyes as compared to healthy contralateral eyes **(Supplementary Figure 2A, B**). Importantly, RGC quantification at 21 days post-IOP elevation depicted similar numbers of preserved RGCs to those of 7 days post-IOP (**Supplementary Figure 2C, D)**.

To evaluate the effect of increasing IOP levels on the integrity of the visual pathway, VEP was measured in each group of animals (**Supplementary Figure 2E**) and response amplitudes were quantified from the peak-to-peak analysis of the first negative component N1 (**Supplementary Figure 2F**). Interestingly, there was a decrease in visual potential amplitude compared to contralateral healthy eye in all experimental groups at day 7 post-IOP. The eyes exposed to 30, 50, and 90 mmHg resulted in reduction of the P1-N1 but the reduction was significant only for IOP >30mmHg. In addition, the magnitude of reduction correlated with the level of applied IOP. VEP signals were not affected by the procedure (**Supplementary Figure 2F**) and VEP amplitudes in IOP-treated eyes 21 days post-IOP were reduced only in groups with IOP elevation >30 mmHg. (**Supplementary Figure 2G**).

To assess potential effects of the IOP elevation on the temporal properties of VEP response, we measured the change in P1 and N1 time-to-peak of the VEP signal. Only the latencies for the 90-mmHg experimental group showed a significant increase for P1 and N1 component, respectively, at 7 days post-procedure. Interestingly this change reverted by day 21 post IOP treatment. None of the other groups exhibited P1 or N1 latency change 7 or 21 days post IOP (**Supplementary Figure 2H, I**).

Next, to explore the extent to which the age of the animal influences the impact of IOP elevation on retinal morphology and function, IOP was unilaterally elevated to 30 mmHg for 1 hour in 3-, 6-, 12-, and 18-month-old wild-type C57BL/6 mice. At day 7 after IOP elevation, animals were sacrificed and immunohistochemistry was performed using anti-Brn3a antibody on flat-mounted retinas. We observed that in 6-month, 12 month and 18-month-old animals RGC count was significantly lower in the retinas exposed to elevated IOP compared to control retinas. The loss of RGCs was drastically higher in 12- and 18-month-old animals compared to 3-month-old animals, suggesting that RGCs in aged animals are more vulnerable to IOP elevation. **(Figure 2A)**.

**Figure 2.**
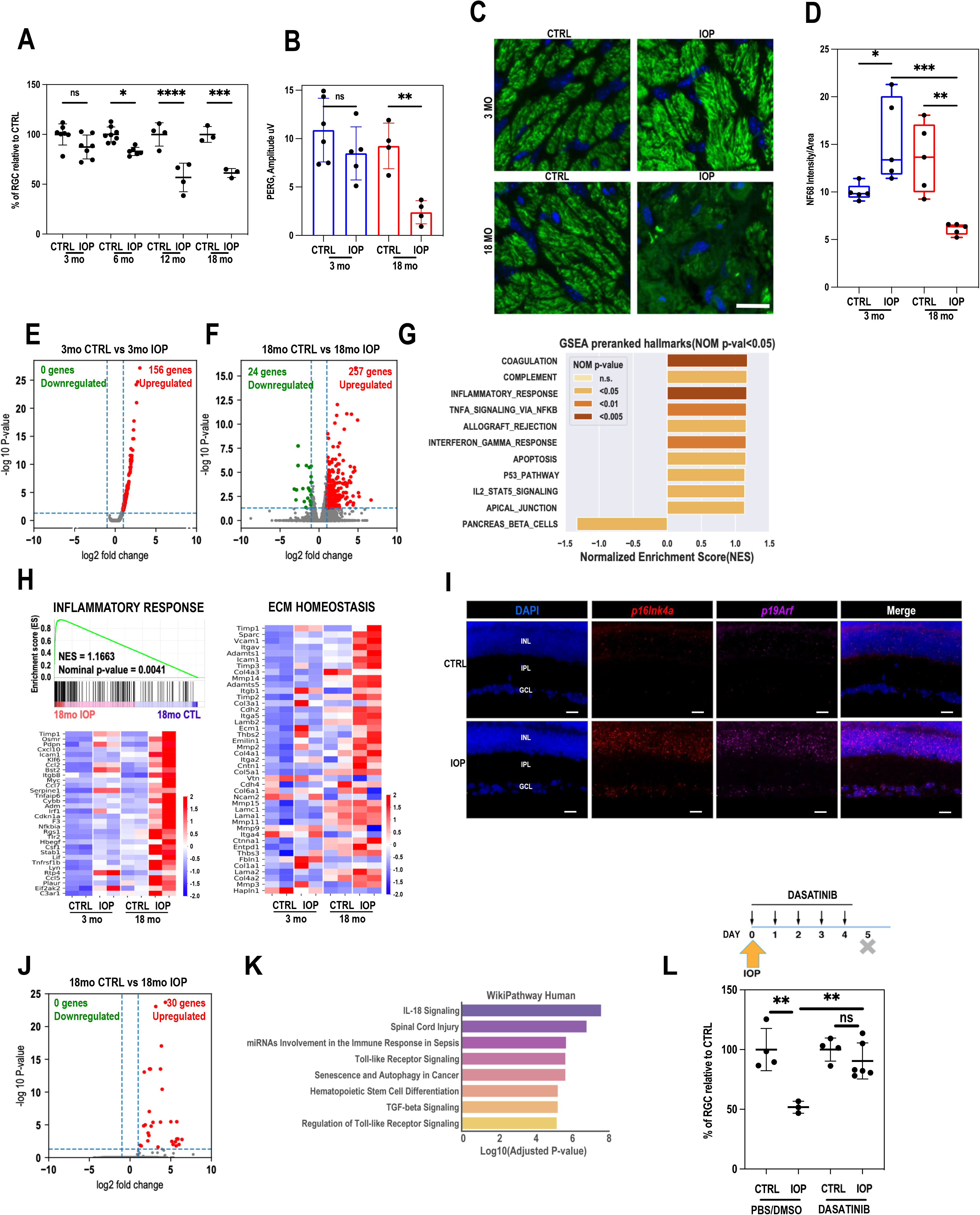
Aged retina is more sensitive to the mild intraocular pressure elevation. **A**. Loss of RGC upon stress is more severe in aged animals. Quantification (Brn3a immunostaining) of RGC loss in 3-month, 6-month, 12-month, and 18-month-old wild type retinas upon 30mmHg IOP on day 5 after treatment (one-way ANOVA, multiple comparisons). **B**. PERG performed on IOP-treated and non-treated (CTRL) eyes on day 5 after IOP treatment in 3-month and 18-month-old C57BL/6J mice. (Multiple comparisons, one-way *P < 0.05; **P < 0.01; ***P < 0.001. **C**. NF68 immunohistochemistry in the glial lamina. (Top-left) Young mice with non-IOP elevation. (top right) Young mice with IOP elevation. (Bottom-left), Aged mice with non-IOP elevation. (bottom right) Aged mice with IOP elevation. **D**. Quantitative analyses of NF68 immunoreactivity (n=5 images/group). Error bars represent SEM. Statistical significance determined using one-way ANOVA. *P < 0.05; **P < 0.01; ***P < 0.001. Scale bar: 100um (all panels). **E-F**. Volcano plots of differentially expressed genes between retinas in 3-month-old (E) and 18-month-old (F) upon 30mmHg IOP. n=2 biological replicates. The numbers of DEGs with defined significance cut-offs were marked in the upper space of the plots. **G**. Gene set enrichment analysis of RNA-Seq data were performed using GSEA-Preranked module. n=2 biological replicates. using the MSigDB hallmark gene set collection (v7.5.1). The bar plot in figure 2G showed all defined significant enriched (nominal pvalue < 5%) hallmark gene sets from pre-ranked GSEA in IOP treated retina compared to the contralateral non-treatment retina from the RNA-seq data were described above. NOM pvalue were shown by a discrete color scale (see scale legend in the figure). **H**. Heatmaps of row z-score of RPKM of each gene in gene sets: Hallmark inflammatory response (left) with its pre-ranked GSEA enrichment plot on top the heatmap and curated senescence pathway genes (right). n = 2 biological replicates. The row z-scores were indicated by a continuous color scale (see scale legend in the figure). **I. J**. Volcano plots showed the transcriptome change immediately after IOP treatment in 18-month retina. n = 3 biological replicates. **K**. Bar graphs showed highly ranked enrichment terms from ontology enrichment analysis of the immediately responded DE genes upon mild IOP using Enrichr. The curated signaling pathway or biological process databases used for the enrichment analysis are WikiPathway (2021 version for human) and MsigDB Hallmark (2020 version). n = 3 biological replicates. **L**. RGC quantification after 5 days of continual Dasatinib treatment. **M**. Histogram plot shows genes expression foldchange of overlapped DEGs upon IOP in both 3-month and 18-month-old mouse retina. **N**. Row z-score heatmap of all up-regulated DEGs immediately after IOP elevation in in 18-month retina. n = 3. O. Gene ontology analysis of the activated DEGs immediately upon IOP in 18-month retina by Enrichr.

To investigate the extent to which age contributes to the loss of RGC function in vision after IOP elevation, IOP was unilaterally raised in 3- and 18-month-old animals and pattern electroretinogram (pERG), measurement that can capture specific response from RGCs, was performed. At 5 days following IOP elevation, not only was the number of RGCs lower in the treated eyes but also 18-month-old animals showed a dramatically reduced PERG amplitude after IOP (mean, 1.97 ± 1.03 μV [SD]; *n* = 3 eyes) in comparison to the control eye (mean, 8.36 ± 1.9 μV; *n*=3 eyes), whereas in young animals no significant difference between treated and nontreated eyes was observed (mean, 6.98 ± 1.78 μV; *n*=3 eyes and mean, 9.92 ± 4.33 μV; *n*=3 eyes, respectively) **(Figure 2B)**.

To further investigate the age-related impact on visual path, we performed immunohistochemistry in the glial lamina cross-sections from 3- and 18-month-old mice treated with IOP elevation (30mmH) using the antibody for neurofilament 68 (NF68), a marker for axons **(Figure 2C)**. The corresponding areas are marked by dashed line (**Supplementary Figure 2J**). In comparison with control young mice with non-IOP elevation, we first found that elevated IOP increased NF68 immunoreactivity in the glial lamina, showing evidence of insulted RGC axons due to IOP elevation. However, in samples isolated from the treated eyeballs from 18-month-old animals, elevated IOP exacerbated axonal degeneration, as observed by a significant reduction of NF68 immunoreactivity in the glial lamina, an effect observed reproducibly in every optic nerve head isolated from the elevated IOP treated eye **(Figure 2D and Supplementary Figure 2J)**. The average region of the axon loss after unique IOP elevation event was assessed to be ∼15% and localized in one sector of the optic nerve **(Supplementary Figure 2J *right*)**.

To capture age-dependent transcriptional changes in the retinas upon unilateral IOP elevation, RNA-seq analysis on bulk retinas collected 2 days after the procedure was performed, when cell loss is not yet observed.^18^ The transcriptome changes were identified using Bioconductor package DESeq2. ^19^ We found that in young retinas, only 156 genes were significantly up-regulated (log2 fold≥1, FDR ≤0.05) and no genes were significantly down-regulated upon IOP stress (log2 fold ≤-1, FDR ≤0.05) **(Figure 2E)**. In contrast, in aged retinas, 257 genes were significantly up-regulated (log2 fold≥1, FDR ≤0.05) and 24 genes were significantly down-regulated upon 30 mm Hg IOP stress (log2 fold ≤-1, FDR ≤0.05) **(Figure 2F)**.

The pre-ranked GSEA showed no significantly enriched pathways in 3-month-old retinas pairs while 9 up-regulated gene sets and 1 down-regulated gene set were significantly enriched at nominal pvalue < 5% between control and IOP treatment in 18-month-old retinas (MsigDB hallmark collection version v7.5.1) **(Figure 2G)**. These gene sets included a series of inflammatory pathways (TNFα signaling via NFκB, INFγ response, complement, IL2-STAT5 signaling and others), p53 and apoptosis pathways and downregulated set of genes recognized as connected with pancreas beta cells pathway. Interestingly, the core genes within significantly enriched pathways (e.g., inflammatory and apoptosis) were robustly upregulated in 18-month-old animals upon stress whereas only slightly upregulated in young retinas **(Figure 2H, Supplementary Figure 2K)**. Similarly, when we plotted heatmaps for senescence and extracellular matrix deregulation pathways^18^ we have observed robust upregulation of the genes in these pathways in 18-month-old IOP-treated retinas but only limited deregulation in 3-month-old-treated retinas. In total, out of 65 top genes dysregulated upon IOP-elevation in 18-month-old retinas, 53 were upregulated more in the old retinas than in the young retinas upon IOP stress **(Supplementary Figure 2M)**. Next, we asked whether the dysregulation of expression of genes detected in bulk RNAseq can be observed in RGCs. For that, 2 days after unilateral 30mmHg IOP elevation, eyeballs were extracted, fixed and cryosections were collected. RNA scope experiment using specific probes for *p16Ink4a* and *p19Arf* cell cycle inhibitors were used. We have observed upregulation of expression of both genes in retinal ganglion cell layer in retinas isolated from IOP-treated eyes **(Figure 2I)**. Surprisingly, both transcripts were also observed in inner nuclear layer in retinas from stressed eyes.

To decipher the general transcriptional program we ran Cytoscape and looked for bundles of pathways that were dysregulated in the aged retina upon stress. Not surprisingly, most of the pathways could be grouped together and named as inflammatory programs **(Figure 2L)**. In addition, we noted bundles related to angiogenesis, apoptosis and finally, phagocytosis.

Next, we sought to discover the pathways that are among the first to be upregulated upon IOP stress in aged retinas. To do so, we collected retinas immediately after the IOP treatment and processed them for RNA extraction and further RNA-seq analysis. Data analysis found 30 genes immediately and significantly upregulated upon stress **(Figure 2J)**. Interestingly, pathways analysis revealed an especially robust immune response upon IOP stress and significant upregulation of senescence and autophagy pathways **(Figure 2K and Supplementary Figure 2M)**. We, therefore, hypothesized that we can use a senolytic approach to remove early senescent cells and potentially protect retinal cells from death. Accordingly, we followed a previously established protocol^18^ **(Figure 2L, *top*)**, and after unilateral 30mmHg IOP treatment of the retina the animals were intraperitoneally injected daily with a senolytic drug, dasatinib. After 5 days, retinas were collected and immunohistochemistry using anti-Brn3a antibody was performed on IOP treated and non-treated flat-mount retinas. Our quantified data show a protective effect of a senolytic drug on RGC number after stress caused by IOP elevation **(Figure 2L)**.

### Age-related epigenetic remodeling upon stress facilitates robust transcriptional response

To understand the chromatin remodeling events that occur after IOP elevation in young and aged retinas, we performed ATAC-seq on retinas isolated three days after the 30 mmHg IOP treatment. Similarly to transcriptional age-related differences, we observed more changes in chromatin accessibility upon IOP stress in aged retinas **(Figure 3A and Supplementary Figure 3A)**. In particular, when we compared the peaks specifically open upon stress, almost four times more regions became opened upon stress in chromatin isolated from 18-months-old retinas than 3-months old retinas. In both cases, most of the regions were either intronic or intergenic **(Figure 3A, B)**. All of these newly opened regions, after cross-referencing with mouse genome database, were located in distant regulatory regions called enhancers. Only very few regions, also localized in enhancers, lost accessibility.

**Figure 3.**
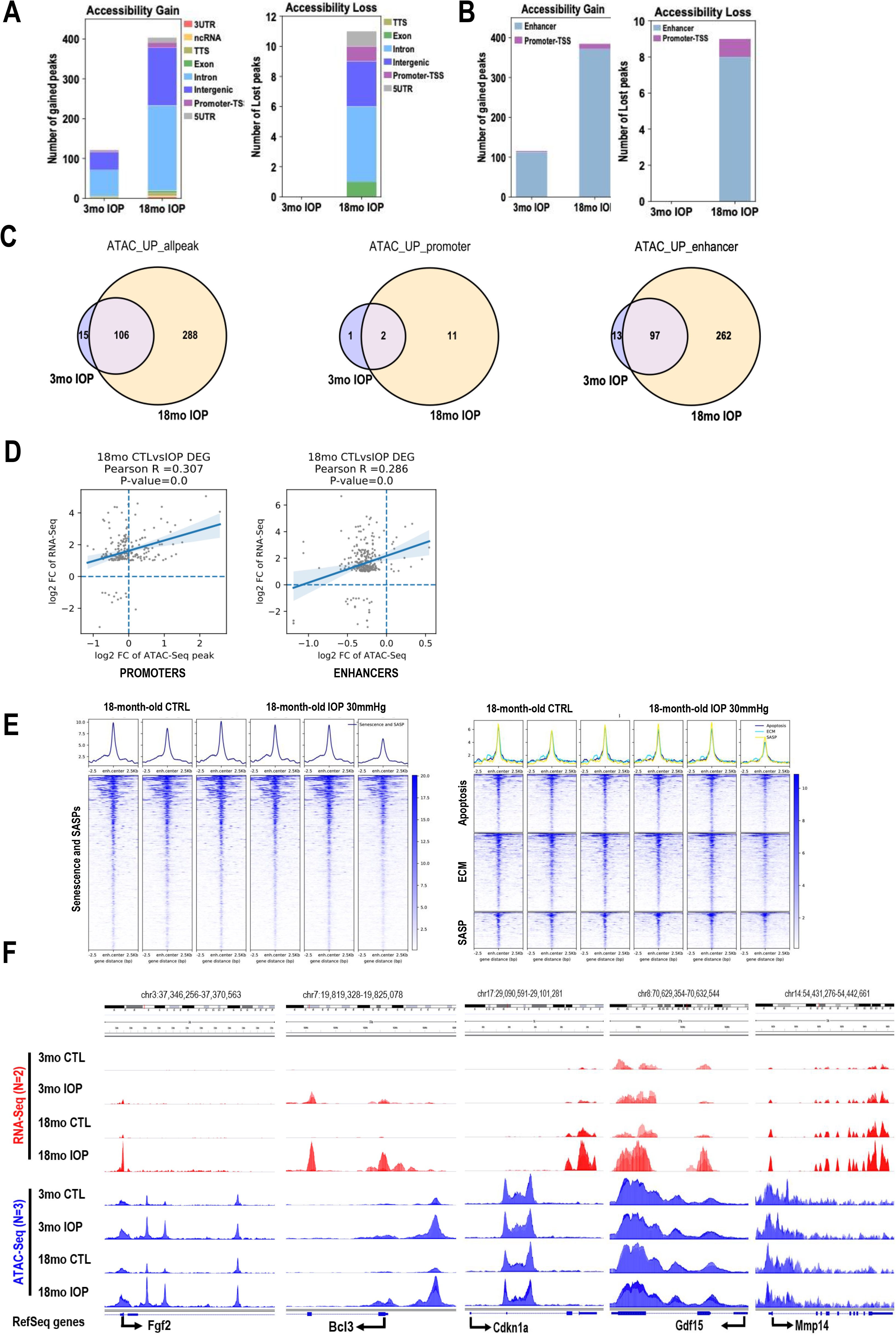
ATAC-Seq analysis of stress-related epigenetic change. **A-B**. Stacked bar plots showed gained or lost accessible regions in both comprehensive distribution (A) and cis-element distribution (B) with cutoffs at 0.05 in FDR and two-fold change of normalized ATAC-Seq signal. Specific genomic regions were color coded. **C**. Venn diagrams showed overlaps of accessibility gain genome-widely or at enhancers/promoters upon mild IOP stress between 3-month-old and 18-month-old retinae. **D**. R-square plots illustrate the correlation between transcription activation and chromatin accessibility changes upon mild IOP treatment in 18-month-old mouse retinae on promoter-TSS regions(left) or Activity-by-contact (ABC) index of both promoters and predicted enhancers with weighted calculation by known *cis*-looping contacts (right). **E**. Profile plots and heatmap plots for chromatin accessibility intensities (deepTools normalized RPKM with bin size 5) over the nearest enhancer regions (2.5Kb flanking distance to center of each region) of curated gene set. In the heatmap plot, higher accessible chromatin regions were shown in darker color (see color legend) and the genomic regions were sorted based on descending order of mean signal value per region in IOP treated 18-month-old mouse retina. n = 3 biological replicates. **F**. The IGV tracks illustrates multiple top-ranked IOP-related DE gene locations in 18-month-old retinae with dynamic changes in chromatin accessibility by dual factors: aging and mild IOP. IGV, Integrative Genomics Viewer.

Next, we looked at the overlap between the epigenetic programs upon stress in 3-months-old and 18-months-old retinas **(Figure 3C)**. Immediately, it became apparent that epigenetic program of young animals almost fully overlaps with the epigenetic program of old animal and only 15 peaks seem to be specific for young animals. We also observed that most stress-induced changes on the chromatin level occurred on enhancers in both 3-month-old and 18-month-old animals, with only 286 additional open enhancers found in aged animal.

Then, statistical scoring based on estimating enhancer activity and enhancer-promoter contact frequency by Activity-by-contact (ABC) model^20^ was performed on ATAC-Seq data which was projected on published Hi-C data in adult mouse retinas^21^ (**Figure 3D**). Each dot in R-square plot represents one activated DE gene with log2foldchange in transcription level versus log2foldchage of accessibility gain at promoter-TSS region(±2.5Kb) (left) or ABC score in the 1Mb flanking range of promoter-TSS (right). We found that there was only a mild correlation (R=0.307) between the presence of open enhancers and gene upregulation, similarly mild correlation (R=286) could be detected on promoters **(Figure 3D)**. Interestingly, when we analyzed pathways of interests, such as senescence, apoptosis, ECM deregulation, we noted no change in enhancer openness for genes in these pathways. In fact, these enhancers were already open in non-treated retina **(Figure 3E)**. Indeed, when we looked at specific examples, we noted that although, some genes highly upregulated specifically in aged retina, only few of them had some changed accessibility in aged retina upon IOP (*Fgf2, Bcl3*), while other genes (*Cdkn1a, Gdf15, Mmp14*) had no change in accessibility in their locus despite significant transcription upregulation **(Figure 3F)**. This intriguing data suggests that other level of regulation (DNA modification, histone modification or other) are involved in final activation of these genes upon signal.

Next, we used HOMER analysis^22,23^ to detect predicted TF binding sites in regions specifically open upon IOP elevation in aged animals **(Supplementary Figure 3A)**. We detected that the top 5 results belonged to STAT family of TFs well known effectors of response to interferons and inflammation.^24^ Interestingly, other TF bindings sites significantly enriched on regulatory elements open upon mild IOP in 18mo retinas were those of homeodomain TFs previously detected as enriched on open elements in aged retina when compared to open elements in young retina (Crx and Otx2, **Supplementary Figure 3B)**. This finding again shows similarities in aging and stress response program.

### Multiple low-level stress occurrences amplify transcriptional response to mild stress and accelerates DNA methylation aging clock

Next, we investigated whether repeated mild IOP elevation in young animal can affect the expression levels of genes involved in the response to stress. In brief, 3-months-old animals were treated with unilateral 30mmHg IOP elevation for 1 hour, four times, every two weeks **(Figure 4A)**. The contralateral eye served as a control. Two days after the final IOP insult, retinas were collected and bulk RNA-seq was performed. Data analysis showed 893 genes significantly up-regulated (log2 fold>=1, FDR<0.05) and 142 genes significantly down-regulated (log2 fold<=-1, FDR<0.05) upon repeated stress, in comparison to nontreated, contralateral eyes **(Figure 4B)**.

**Figure 4.**
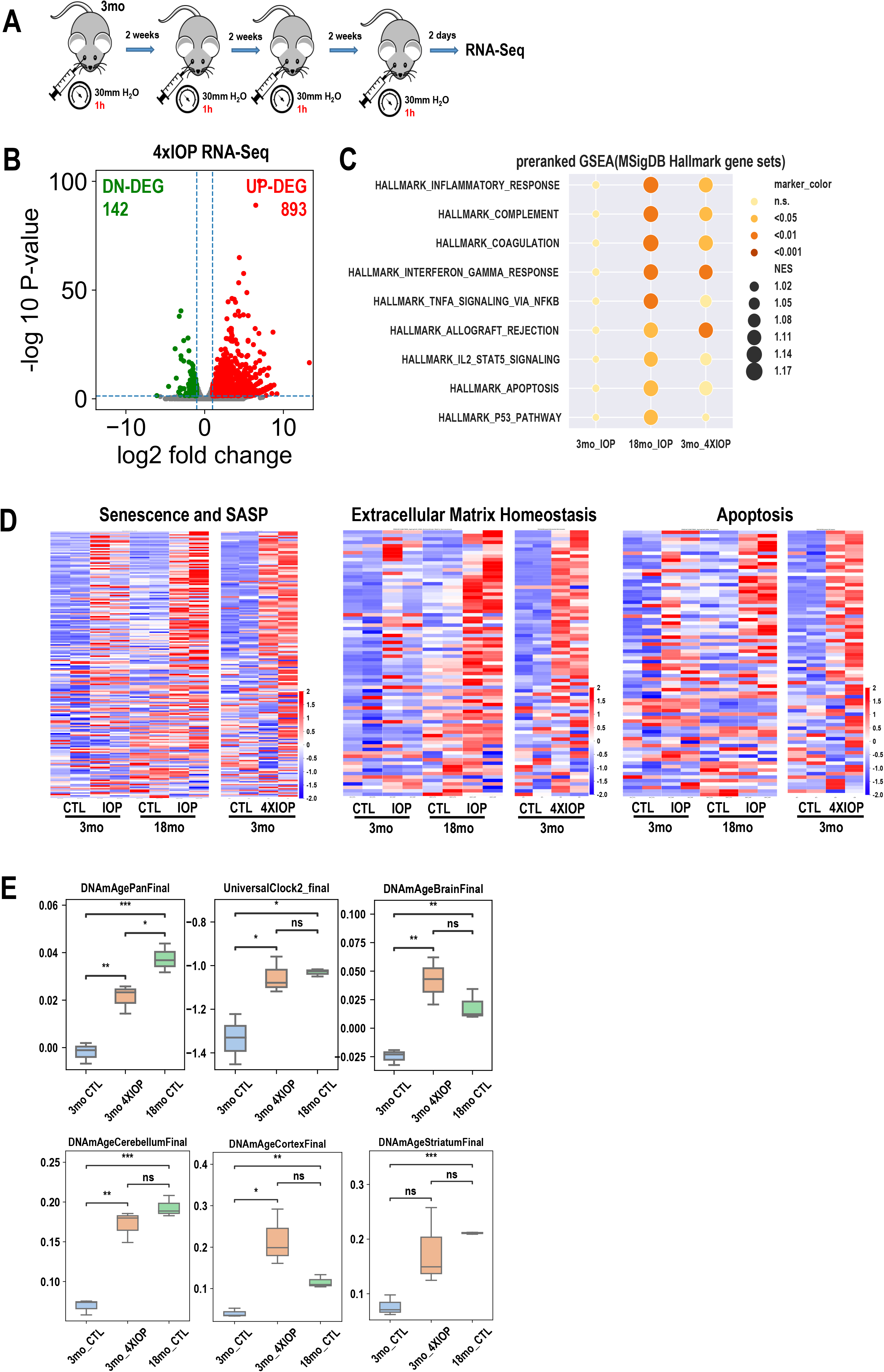
Recurring stress accelerates aging of the retina. **A**. Scheme of the experiment. 3-months-old mice were exposed to unilateral IOP elevation (30 mmHg) for 1 hour. The experiment was repeated every two weeks over 2-month period, in total 4 unilateral (OD) IOP elevation experiments per animal. The contralateral eye served as a control. Two days after the final IOP, retinas were collected and bulk RNA-seq was performed. **B**. Volcano plots showed the transcriptome change upon multiple times of IOP treatment. n = 2 biological replicates. **C**. Dot plot show gene expression regulation pattern by preranked GSEA enrichment analysis of one-time IOP treatment on 3-month and 18-month retinae and multiple IOP treatment on 3-month retinae. n = 2 biological replicates. The NOM p-value of each term is shown by a discrete color scale of the dot. Normalized enrichment score (NES) is shown by dot size (see figure legend). C Row Z-score Heatmap of gene expression of curated gene pathways. **D**. Predicted age (by unit of years) by aging clock microarray analysis of one-time IOP treated 3-month and 18-month retinae and multiple IOP treated 3-month retinae. n = 3 biological replicates.

The pre-ranked GSEA revealed several pathways involved in inflammatory response, apoptosis, and DNA repair to be significantly upregulated. Interestingly, the list of significantly upregulated pathways closely resembled that of pathways upregulated in 18-month-old retinas upon 30mmHg IOP elevation **(Figure 4C)**. Moreover, when we compared pathways such as senescence and SASPs, ECM deregulation and apoptosis, the pattern of gene expression in 4xIOP retinas resembled the expression pattern of aged retina after IOP and not that of young retina after the stress **(Figure 4D)**. This suggests that recurrent stress on young tissue can induce changes in its ability to respond to stress making it transcriptionally similar to aged retina.

Following this finding, we asked whether repeated IOP elevation in young retinas can induce age-related changes that could be measured by an independent method to that of gene expression. We therefore investigated whether our treatment induced changes in the DNA methylation pattern that could be quantified to assess the “aging clock” through the pattern of DNA methylation, DNA isolated from the control and 4xIOP retinas. DNA samples including those isolated from 18-month-old mouse retinas were sent to the Epigenetic Clock Development Foundation that subsequently performed blinded DNA methylation measurement and analysis. The Illumina Horvath Mammalian Methylation 320k Chips designed for mouse methylation studies^25^ were used and testing was performed for the pan-tissue mouse clock as well as for liver, blood, brain, muscle, heart, cortex, striatum, cerebellum, tail, kidney, skin, and fibroblasts. Since there is no retina specific DNA methylation clock, we developed a strategy to evaluate the differences between the tissues based on the clocks included in the test. First, we plotted the data generated using all pan-tissue and universal clocks included in analysis **(Figure 4E)**. We noted that all of them detect not only the difference in DNA methylation age (DNAmAge) between 3-months-old and 18month-old tissues but also show that 3-month-old tissue after 4xIOP exhibits DNA methylation levels significantly higher than control retina. Furthermore, the detected “DNA methylation age” in repeatedly stressed 3-month-old tissue is close to that of 18-months-old retina. Next, we plotted the data generated from methylation patterns of different regions in the central nervous system other than retina – cortex, cerebellum, striatum, and whole brain **(Figure 4E, Supplementary Figure 4C,D)**. This analysis again detected differences in the methylation clock between 3-month-old and 18-month-old retinas, and most importantly between nontreated and 4xIOP treated retinas, suggesting that repeated mild stress is able to change the methylation pattern of DNA in stressed tissue. Finally, we plotted the data using the clocks generated on the DNA methylation data from tissues unrelated to the retina – blood and fibroblasts **(Supplementary Figure 4E)**. In contrast to CNS related clocks, these clocks were unable to reproducibly detect a difference in age between young and old tissue and have shown little of consistency of the age difference between untreated vs. treated young retinas.

## DISCUSSION

In this work we focused on understanding differences in stress response between young and old animals in an experimental model of ocular hypertension. First, we describe a significant decline in visual function with age. These changes correlate with transcriptional changes in the retina, showing increase of inflammatory and cell death pathways, as well as degradation of ECM integrity during aging. Our epigenetic analysis has shown a significant increase in chromatin accessibility with age, mostly on enhancers. Next, we show that upon mild IOP elevation, aged retina further activates inflammatory, senescence and other age-related pathways and that these changes occur also in RGCs. Interestingly, the response to the same, mild stress, in young tissue is much weaker and does not trigger RGC death. Finally, we show that we can induce very strong stress response in young tissue upon mild hypertension once the retina is “primed” with previous instances of mild IOP elevation. We conclude that several occurrences of stress in young retina accelerate tissue age as estimated by transcription profile and by DNA methylation clock.

Glaucoma is characterized by slow, progressive loss of RGCs, sectorial progression in optic nerve degeneration and eventual vision loss^.26,27^ Here, we characterize new animal model relevant to the human disease that can allow study of the time course of molecular events in the retina upon mild ocular hypertension. We show that 18-month-old animals loose RGCs and vision as measured by PERG upon 30mmHg IOP elevation for 1h, while 3-month-old animals do not show these phenotypes. In addition, cross-section of optic nerve of the treated and non-treated eyes shows sectorial loss of axons but only in old animals. Finally, our molecular analysis detects senescence as one of the pathways involved in the response to stress mechanism of the retina upon IOP elevation, what we previously documented in human glaucomatous tissue.^28^

A main hurdle in developing new therapies and understanding the molecular mechanism of glaucoma is the lack of an animal model that recapitulates all aspects of the disease. While some laboratories focus on developing animal models of chronic IOP elevation^29^,^30^, others use mice with natural mutations that slowly develop age-related hypertension and RGC loss. ^31^,^32^,^33^ We have been particularly interested in studying an animal model that allow us to capture the time course of molecular changes upon glaucomatous stress, such as mild ocular hypertension. Since age is the strongest risk factor for developing glaucoma, we developed and characterized the aged animal model for our future molecular studies.

The molecular analysis of transcriptional changes between IOP treated and non-treated retinas have detected a high number of pathways involved in innate immunity and sterile inflammation. The significant changes can be observed in aged animals, however most of the same genes are also slightly upregulated in 3-month-old animals without reaching significance. Interestingly, a similar coordinated inflammatory program was observed when we compared transcriptional differences between young and old retina transcriptomes suggesting that IOP stress was able to elicit age-related inflammatory response in the tissue. In fact, the list of pathways upregulated in aged and aged-IOP-treated retinas is similar, and analogous to the set of pathways called “inflammaging” in previous studies.^3^ Since RGCs degenerate early in glaucoma, we sought to find whether observed transcriptional changes happen in RGCs, as well. We observed that upon 30mmHg IOP elevation, RGCs express hallmarks of senescence: the cell cycle inhibitors *p16Ink4a* and *p19Arf* (**Figure 2I**),) as we have previously shown in human glaucomatous retinas. These observations further confirm the ability of neurons, postmitotic cells, to assume senescent phenotype^11,34,35^.

We then describe the pathways induced immediately upon IOP elevation in aged retinas. We have isolated retinas immediately after the procedure and compared treated and non-treated retina transcriptomes. Only 30 genes were upregulated upon IOP elevation, and none was downregulated **(Figure 2J)**. To our surprise, several immune response pathways were significantly enriched in this small group of transcripts. Interestingly, these were among the first pathways in which we noticed senescence as being significantly upregulated **(Figure 2K)**. Moreover, we show, even in aged retina, that RGCs can be protected from ocular hypertension-related death with senolytic drugs that remove senescent cells. These data suggest that senescence and some set of inflammatory genes are pre-programmed to swiftly respond to stress in aged retina. This is why we asked whether changes in DNA accessibility can explain the different levels of response to the same signal between young and old tissue.

The chromatin accessibility assay data is puzzling. During the 15 months of retina aging only ∼250 sites in the chromatin opened, most of them being enhancers (**Figure 1H**). Even fewer sites lost their accessibility (∼70). Given the fact that one gene can have several regulatory elements and there are ∼4000 deregulated transcripts in aging (**Figure 1D)**, we assume that most of the regulation of the expression of the age-related changed transcripts is not related to the level of DNA accessibility. In the mild IOP data sets (RNAseq vs. ATAC-seq), the numbers of transcripts are similar to numbers of detected open chromatin sites, however we could not detect correlation of upregulation of expression and increase in enhancer accessibility in these samples (**Figure 3**) suggesting other pathways being involved in regulation of activation of expression of these enhancers. Further studies are needed to address this point.

After looking at the conserved TF binding sites on open regions, we have detected that STAT factors were particularly highly represented, in agreement with our transcriptomic data (**Supplementary Figure 3B**). Interestingly, on the list of the TF binding sites on open regions in stressed young and aged retinas, we noticed binding sites of homeodomain TFs, such as Crx and Otx2, detected previously as enriched in sites opened in natural aged retina **(Figure 1I)**. This observation further shows similarities between the stress response and aging.

Following these findings, we asked whether repetitive mild stress in young animals can induce accumulation of stress response and tissue damage. We allowed two weeks of recovery between each 30mmHg IOP elevation **(Figure 4A)**. Transcriptomic data was clear – retinas that underwent recurrent mild stress exhibited significant upregulation of almost 900 genes **(Figure 4B)**. GSEA analysis indicated high upregulation of inflammatory pathways and deregulation of apical junction **(Supplementary Figure 4A**). Metascape analysis clustered all pathways in inflammation-related bundles, highlighting the level of sterile inflammation induced upon repeated stress (**Supplementary Figure 4B)**. The response to recurrent stress was significantly higher than in the 3-month-old retina treated one time. In fact, the transcriptional response of the 4xIOP treated retina was very similar to that of the 18-month-old retina upon a single 30mmHg IOP elevation **(Figure 4D)**. Therefore the transcriptional phenotype suggested that the tissue aged with several rounds of stress.

DNA methylation-based clocks were developed around 10 years ago, to objectively and independently, assess the chronological age of the human, animal, tissue, and any nucleated cells.^36,37,38,39^ As DNA methylation levels are strongly correlated with age, the profile of genome wide CpG methylation status can be used to estimate the age of a specimen. Interestingly, it has been shown that although similar, each tissue has its own pace of DNA methylation change and, therefore, one can develop a specific clock for any given tissue.^40,41^ These “DNA methylation clocks” can be then used to follow the impact of diseases or environmental cues on aging. They has been shown to be especially useful in population genetic studies by uncovering new associations between aging and different health conditions.^39,42,43^ As there is no mouse retina specific clock, we decided to work with clocks that were developed to estimate age of all tissues and/or all animals (e.g. DNAmAgePanFinal, UniversalClock_final) developed by the Horvath laboratory.^44^ Importantly, due to the lack of a retina specific clock, the numbers of years (age of tissue) obtained in the assay should be treated as relative levels. Our DNA methylation clock data were quite remarkable. The PanTissue and Universal clocks showed that 3-month-old and 3-month-old 4xIOP tissues were no longer the same epigenetic age. Moreover, 4xIOP tissue was now closer, epigenetically, to the 18-month-old DNA methylation age. We also looked at the clocks developed for several regions of CNS as well as the whole brain. In all cases, 3mo animals were correctly separated from 18mo animals, indicating the difference in age. But also using CNS specific clocks, our data indicated that repetitive mild stress on young tissue induced accelerated “ticking” of the DNA methylation clock and also showed that 4xIOP tissue was older than its contralateral control. These transcriptomic and DNA methylation data collectively imply that multiple instances of mild IOP can accelerate aging of the retina.

In humans, IOP has a circadian rhythm.^45,46^ In healthy individuals, it oscillates typically in the 12-21 mmHg range and tends to be highest in approximately two thirds of individuals during the nocturnal period.^45,47^ It has been suggested that dysregulation of the diurnal IOP cycle, with higher daily amplitudes and longer periods of ocular hypertension, might be the cause of some cases of glaucomatous optic neuropathy.^45,48^ Since IOP at a level sufficient to cause ischemia is significantly more severe than even the highest circadian fluctuations in glaucoma patients, it is critical to study glaucoma-related phenomena in the models more closely recapitulating human disease. In this study, we have shown that even moderate hydrostatic IOP elevation to 30 mmHg for 1 hour, a level relevant to human disease, results in RGC loss and corresponding visual defects.

In sum, we have shown that susceptibility to stress changes with age and is pre-conditioned epigenetically. One of the earliest pathways activated upon IOP elevation in 18-month-retina is senescence. Moreover, the use of a senolytic drug can protect RGCs from death. Finally, we have shown that multiple instances of stress in young animal may cause changes in transcriptional and DNA methylation patterns indicating accelerated aging. These results emphasize the importance of early diagnosis and prevention as well as age-specific management of age-related eye-diseases, including glaucoma.

## MATERIALS AND METHODS

### Hydrostatic intraocular pressure (IOP) elevation

Animals were anesthetized with an intraperitoneal injection of ketamine/xylazine cocktail, (100 mg/kg and 10 mg/kg, respectively), their eyes numbed with a drop of proparacaine (0.5 %, Bausch-Lomb) and dilated with a drop of tropicamide (1 %, Alcon Laboratories) followed by a drop of phenylephrine (2.5%, Akorn Pharmaceuticals, Lake Forest, IL). To achieve IOP elevation, a 33-gauge needle was advanced trans corneally into the anterior chamber under visual control. Elevation of IOP was achieved by instilling the aqueous chamber of the eye with Balanced Salt Solution (Alcon Laboratories, Fort Worth, TX) through an IV infusion set. The desired level of IOP increase was achieved by pumping infusion bag under control of sphygmanometer. Stable elevated IOP was maintained for 60 minutes, controlled by IOP measurements using a veterinary rebound tonometer (Tonovet). Both eyes were lubricated with an ophthalmic lubricant gel (Alcon Laboratories) during the protocol. Following the procedure one drop of topical antibiotic was applied to the treated eye (Ofloxacin 0.3%, Apexa). The anesthesia was reversed with Atipamezole (0.1 mg/kg) and animals recovered on a isothermal pad until awake. The contralateral eye without IOP elevation served as a healthy non-IOP control (CTRL). Animals were treated at 3 pressure levels (30, 50, and 90 mmHg), and studied at two timepoints (7 and 21 days).

To control for procedural effects, two additional pressure level cohorts – 7 mmHg, and 15 mmHg – were investigated at the day 7 time point. The 7 mmHg eyes were cannulated without elevating pressure through the IV infusion set. Each experimental group consisted of n = 6 animals (3 male, 3 female). Additionally, we studied a cohort of 6-month-old mice (3 male, 3 female) 7 days after following unilateral hypertension to 30 mmHg for 1 h.

### Neurophysiology (Figure 1)

Animals were initially anesthetized with 2% isoflurane in a mixture of N_2_O/O_2_ (70%/30%) then placed into a stereotaxic apparatus. A small, custom-made plastic chamber was secured to the exposed skull using dental acrylic. After one day of recovery, re-anesthetized animals were placed in a custom-made hammock, maintained under isoflurane anesthesia (1-2% in N_2_O/O_2_), a craniotomy was performed, and multiple single tungsten electrodes were inserted into V1 layers II-VI. Following electrode placement, the chamber was filled with sterile agar and sealed with sterile bone wax. Animals were then sedated with chlorprothixene hydrochloride (1 mg/kg; IM; ^49^) and kept under light isoflurane anesthesia (0.2 – 0.4% in 30% O_2_) throughout the recording procedure. EEG and EKG were monitored throughout, and body temperature was maintained with a heating pad (Harvard Apparatus, Holliston, MA).

Data was acquired using a multi-channel Scout recording system (Ripple, UT, USA). Local field potentials (LFP) from multiple locations at matching cortical depths were band-pass filtered from 0.1 Hz to 250 Hz and stored along with spiking data at a 1 kHz sampling rate. LFP signal was aligned to stimulus time stamps and averaged across trials for each recording depth in order to calculate visually evoked potentials (VEP)^50–52^. Single neuron spike signals were band-pass filtered from 500 Hz to 7 kHz and stored at a 30 kHz sampling frequency. Spikes were sorted online in Trellis (Ripple, UT, USA) while performing visual stimulation. Visual stimuli were generated in Matlab (Mathworks, USA) using Psychophysics Toolbox ^53–55^ and displayed on a gamma-corrected LCD monitor (55 inches, 60 Hz; 1920 × 1080 pixels; 52 cd/m^2^ mean luminance). Stimulus onset times were corrected for monitor delay using an in-house designed photodiode system^56^. Visual responses were assessed according to previously published methods ^51,56,57^. For recordings of visually evoked responses, animals were tested with 100 repetitions of a 500 ms bright flash of light (105 cd/m^2^).

### Local Field Potential (LFP) Analysis

Amplitude of response was calculated as a difference between the peak of the positive and negative components of the VEP. Response latency was defined as the time from stimulus onset to maximum response. Maximum of the response was defined at the larger of the negative or positive peak.

### Visual evoked potential (VEP) (Figure 2 and Supplementary Figure 2)

VEP measurements were recorded at 7 and 21 days post-IOP elevation according to a previously published protocol. Mice were dark adapted for at least 12 hours before the procedure. Animals were anesthetized, their eyes dilated as in the IOP elevation protocol. The top of the mouse’s head was cleaned with an antiseptic solution. A scalpel was used to incise the scalp skin, and a metal electrode was inserted into the primary visual cortex through the skull, 0.8 mm deep from cranial surface, 2.3 mm lateral to the lambda. Platinum subdermal needle (Grass Telefactor) was inserted through the animal’s mouth as reference, and through the tail as ground. The measurements commenced when baseline waveform became stable, 10-15 s after attaching the electrodes. Flashes of light at 2 log cd.s/m^2^ were delivered through a full-field Ganzfeld bowl at 2 Hz. Signal was amplified, digitally processed by the software (Veris Instruments), then exported and peak-to-peak responses were analyzed in Excel (Microsoft). To isolate VEP of the measured eye from the crossed signal originating in the contralateral eye, black aluminum foil eyepatch was used to cover the eye not undergoing measurement. For each eye, peak-to-peak response amplitude of the major component N1 in IOP eyes was compared to that of their contralateral NIOP controls. Following the readings, the animals were euthanized, their eyes collected and processed for immunohistochemical analysis.

### Optomotor responses

Optomotor responses (OMRs) were recorded using commercially available qOMR setup. (PhenoSys GmbH, Berlin, Germany). The software automatically tracks animal head movement in relation to the moving grating stimulus and calculates correct/incorrect tracking behavior presented as quantitative optomotor index (qOMR). The mouse was placed on elevated platform in OMR arena. Grating stimuli (rotating at 12°/s) were presented for ∼12 minutes per trial. The spatial frequency of the grating was set at 0.2 cycles per degree of visual angle. The stimulus was presented at differing contrasts between the light and dark: 3, 6, 10, 12.5, 15, 17.5, 20, 25, 37.5, 50, 75, 100 contrast. Stimulation at each contrast level lasted 60 s. Each mouse was tested in 4 photopic trials and 4 scotopic trials. Each mouse was firstly tested in photopic conditions and then in scotopic. For the scotopic part of the experiment (resembles nighttime light levels) the animals were dark-adapted for 12h before the experiments. The OMR arena was dimmed using density 4 filters in front of the stimulus displays. The luminance for scotopic part was ∼0.03 lux. Results for 100% contrast were excluded due to adjustment (mice adjustment to the system). qOMR results that yielded correct/incorrect ratio lower than 0.8 were excluded. Results from the trials were averaged for analysis of each individual mouse. Each data point represents average of all mice tested. GraphPad prism was used for statistical analysis and to generate curves.

### Pattern electroretinography

All mice were dark-adapted for 12h before pattern electroretinography (PERG) was recorded. PERG recordings were obtained with the commercially available Celeris ERG platform equipped with pattern ERG stimulators (Diagnosys LLC, USA). Prior to experiment all mice where anesthetized with Ketamine/Xylazine (100 mg/kg and 10 mg/kg, respectively) injected intraperitoneally.

The eyes were anesthetized with proparacaine (0.5 %, Bausch-Lomb), next the eyes were dilated using phenylephrine (2.5%, Akorn Pharmaceuticals, Lake Forest, IL) and tropicamide (1 %, Alcon Laboratories) commercially available eyedrops. The eyes were lubricated with corneal gel. After anesthesia and eye dilation pattern erg stimulators were placed on corneal surface. For all mice 400 sweeps (reads) per eye were recorded. PERG amplitude was measured between first positive peak (P1) and second negative peak (N2).

### Senolytic drug treatment

The experimental group of animals was treated by intraperitoneal (IP) administration of dasatinib (Sigma, 5 mg/kg) after IOP elevation (see below), and a control group of mice was sham-treated vehicle (PBS/DMSO). Each mouse underwent unilateral hydrostatic pressure-induced IOP elevation to 30 mm Hg, with the contralateral eye left as an untreated control. The mice were IP injected intraperitoneally with dasatinib at day 0 (IOP elevation day) and continued for four consecutive days **(Figure 2L)**. At day 5, animals were euthanized, and retinas were isolated and immunostained with anti-Brn3a antibody to evaluate the number of RGCs. All drugs were prepared according to the UC Irvine Institutional Animal Care and Use Committee (IACUC) standards. To ensure a sterile environment, compounds were prepared under the tissue culture hood using sterile PBS. The final solution was filtered through a 0.22-μm PES membrane just before injection. Tips, tubes, and syringes were sterile.

### Immunohistochemistry for RGC count (Anti-Brn3a and Anti-RBPMS)

Eyes were fixed in 4% paraformaldehyde (PFA) in PBS for 2 hrs then transferred to PBS. The retinas were extracted through dissection and were flat-mounted on microscope slides and blocked for 1 hr in blocking solution (10% bovine serum albumin (BSA) diluted in 0.5% Triton-X/PBS). Standard sandwich assays were performed to immunostained using anti-Brn3a antibodies (Millipore, MAB1585) and anti-RBPMS antibodies following secondary AlexaFluor 555 anti-mouse (Invitrogen, A32727) and secondary AlexaFluor 647 anti-rabbit (Invitrogen, A21245). The samples were then mounted (ProLong Gold antifade reagent, Invitrogen P36934) and imaged using a fluorescent microscope at 20x magnification (Biorevo BZ-X700, Keyence). Peripheral retina regions were chosen to optimize the quality of the micrographs. The micrographs, which were 0.39 mm^2^ per region, were split into 4 equal parts to exclude disqualified, damaged regions. Labeled cells were manually quantified with the “cell counter” function of the software Fiji.

### Optic nerve head - tissue preparation and immunohistochemistry

Mice were anesthetized by an IP injection of a cocktail of ketamine/xylazine as described above prior cervical dislocation. For immunohistochemistry, the optic nerve head (ONH) tissues were dissected from the eyeballs and fixed with 4% paraformaldehyde (Sigma) in phosphate buffered saline (PBS, pH 7.4, Sigma) for 2 h at 4 °C. ONHs were washed several times with PBS then dehydrated through graded levels of ethanol and embedded in polyester wax.

Immunohistochemical staining of 7 µm polyester wax sections of full thickness retina were performed. Sections from polyester wax blocks from each group (*n* = 3 ONHs/group) were used for immunohistochemical analysis. To prevent non-specific background, tissues were incubated in 1% bovine serum albumin (BSA, Sigma)/PBS for 1 h at room temperature before incubation with mouse monoclonal NF68 antibody for 16 h at 4°C. After several wash steps, the tissues were incubated with the Alexa Fluor-488 conjugated donkey ant-mouse IgG antibody for 4 h at 4°C and subsequently washed with PBS. The sections were counterstained with the nucleic acid stain Hoechst 33342 (1 µg/ml; Invitrogen) in PBS. Images were acquired with Keyence All-in-One Fluorescence microscopy (BZ-X810, Keyence Corp. of America, IL, USA). NF68 protein fluorescent integrated intensity in pixel per area was measured using the ImageJ software. All imaging parameters remained the same and were corrected with the background subtraction.

### Cell dissociation and sorting

For one sample of selected RGCs two dissected retinas were dissociated as previously described ^58,11^ and stained with CD90.2(Thy-1.2)-FITC antibody (Thermo Fisher REF#11-0902-81). Briefly, dissociated cells from two retinas were resuspended in 100ul of staining buffer (DPBS with 2% FBS(v/v) and 10mM EDTA) containing 1:50 dilution of Thy1.2-FITC antibody and incubated for 30min in complete darkness. After incubation cells were washed in enriched PBS (1mg/mL D-glucose, 4% BSA(w/v) and 0.5mg/mL insulin in DPBS (Ca^2+^,Mg^2+^)) and prepared for sorting using microfluidic cell sorting (WOLF® Cell Sorter NanoCellect) were FITC was used as reported for selection. Enriched PBS was used as both dilution and sheath buffer during sorting. Approximately 100,000 cells were isolated in each experiment and processed for RNA extraction.

### RNA-Sequencing (Next Generation Sequencing)

Fresh retina tissue of each mouse eye was homogenized thoroughly in TRIzol Reagent (Cat no.15596026, Invitrogen) followed by RNA isolation. For pellet of immunoselected RGCs – cells were resuspended in Trizol reagent. The polyadenylated (poly-A) transcripts from the isolated total RNA sample were enriched and performed strand specific mRNA-Seq libraries preparation by TruSeq kit followed by Next Generation Sequencing on NovaSeq 6000 System (Flow Cell Type S4). mRNA-Seq raw reads data was mapped to mm10 assembly mouse genome by homer (v4.11) aligner tool STAR. The uniquely mapped tags in exons on minus strand (one isoform per locus) were quantified by analyzeRepeats.pl. Raw reads count was normalized to experiment totals and performed differential expression based on the negative binomial distribution by DESeq2 v1.24.0 program^19^ with FDR q-value significance cut-off at 0.05 and log2(fold change) significance cut-off at 1 for up-regulation or −1 for down-regulation.

### Gene set enrichment analysis (GSEA)

The Gene Set Enrichment Analysis (GSEA v4.2.3) was performed for pairwise differential expression comparisons between two phenotype groups of RNA-Seq samples in the pre-ranked module.^59,60^ The target chip annotation platform was Mouse_RefSeq_Accession_Extended_Human_Orthologs_MSigDB.v7.5.1. chip and gene set collection applied was the condensed hallmark pathways (v7.5.1) from the Molecular Signatures Database (MSigDB).^16,61^ The differential expression metrics were ranked by Rank Metric Score (Log2Fold Change*Log10(FDR q-value). Enrichment analysis was performed using weighted (p=1) scoring calculation scheme with 1000 permutations. Gene sets with size larger than 500 and smaller than 5 were excluded from the analysis.

### Metascape gene enrichment and functional analysis

List of RefSeq identifiers of differential expressed (DE) genes (FDR q-value ≤ 0.05; log2foldchange ≥ 1 as significant up-regulated DE genes or ≤-1 for downregulated) from transcriptome analysis of each comparison pair^17^ were abundantly annotated by retrieving from multiple databases. The given list of annotated gene identifiers was then statistically enriched into terms by selected ontology sources which contain Gene Ontology Biological Processes, KEGG Pathways, Reactome Gene Sets and WikiPathways. Terms with a p-value < 0.01, a minimum count of 3, and an enrichment factor > 1.5 (the enrichment factor is the ratio between the observed counts and the counts expected by chance) are collected and grouped into clusters based on their membership similarities. To further capture the relationships among the terms, the subsets of the enriched terms were rendered as interaction networks where terms with a similarity > 0.3 are connected by edges and formed intra- and inter-clusters The Metascape enrichment networks were finally visualized in nodes-and-edges by Cytoscape (v3.9.0). Node size is proportional to the number of input genes in the term. Each node represents an enriched term and was either colored by cluster ID or colored in discrete scale by FDR q-value.

### ATAC-Sequencing

Fresh retina tissue of each mouse eye was re-suspended immediately in 1 ml ice cold nuclei permeabilization buffer (5%BSA, 0.2% (m/v) NP40, 1mM DTT in PBS solution) with 1X complete EDTA-free protease inhibitor and homogenized using a syringe and needle followed by slow rotation for 10 minutes at 4°C. The nuclei suspension was then filtered through a 40uM cell strainer and centrifuged for 5 minutes at 500xg at 4°C. The nuclei pellet was resuspended in an ice cold 50 µl tagmentation buffer. Nuclei concentration was adjusted to 2,000-5,000 nuclei/µl and 10 µl of the suspension was used for tagmentation. 0.5 µl Tagment DNA Enzyme 1 (FC-121-1030, Illumina) was added to the 10uL suspension. The reaction mix was thoroughly pipetted and incubated 30 minutes with 500 rpm at 37 °C. After the tagmentation reaction completed, the DNA was isolated using Qiagen PCR Purification Kit (Cat.#.28304, Qiagen) and elute in 20 uL Elution Buffer. The eluted DNA fragments were then amplified by PCR with Nextera compatible indexed sequencing i5 and i7 adapters using NEBNext 2x PCR Master Mix PCR kit (M0541, NEB). The amplified DNA library was fragment size selected from 200bp to 800bp using Ampure XP beads (A63880, Beckman Coulter). The quality of the ATAC-Seq libraries was assessed by Agilent 2100 bioanalyzer (Agilent Technologies, Inc.). ATAC-Seq libraries was pooled and run on a NovaSeq 6000 System (Flow Cell Type S4) Illumina sequencer with a paired-end read of 100 bp to harvest about 50 million paired-end reads per sample.

### DNA methylation clocks analysis

The Illumina Horvath Mammalian Methylation 320k Chips designed for mouse methylation studies^25^ were used. This enables the quantitative interrogation of more than 320,000 CpGs per sample, including >285,000 specific to mice and 37,000 conserved mammalian loci. 250ng DNA of each sample (n=4 for each group) underwent DNA extraction, bisulfite conversion, DNA methylation arrays, and statistical analysis as previously described^25,62^. Testing was performed for the pan-tissue mouse clock as well as for liver, blood, brain, muscle, heart, cortex, striatum, cerebellum, tail, kidney, skin, and fibroblasts.

## CONFLICT OF INTEREST

None

## FUNDING

Work in Dorota Skowronska-Krawczyk laboratory is funded by R01 EY027011. The authors would also like to acknowledge an unrestricted RPB grant to the University of California, Irvine, Department of Ophthalmology. The International Centre for Translational Eye Research (MAB/2019/12) project is carried out within the International Research Agendas program of the

Foundation for Polish Science co-financed by the European Union under the European Regional Development Fund.

## FIGURE LEGENDS

**Supplementary Figure 1.**
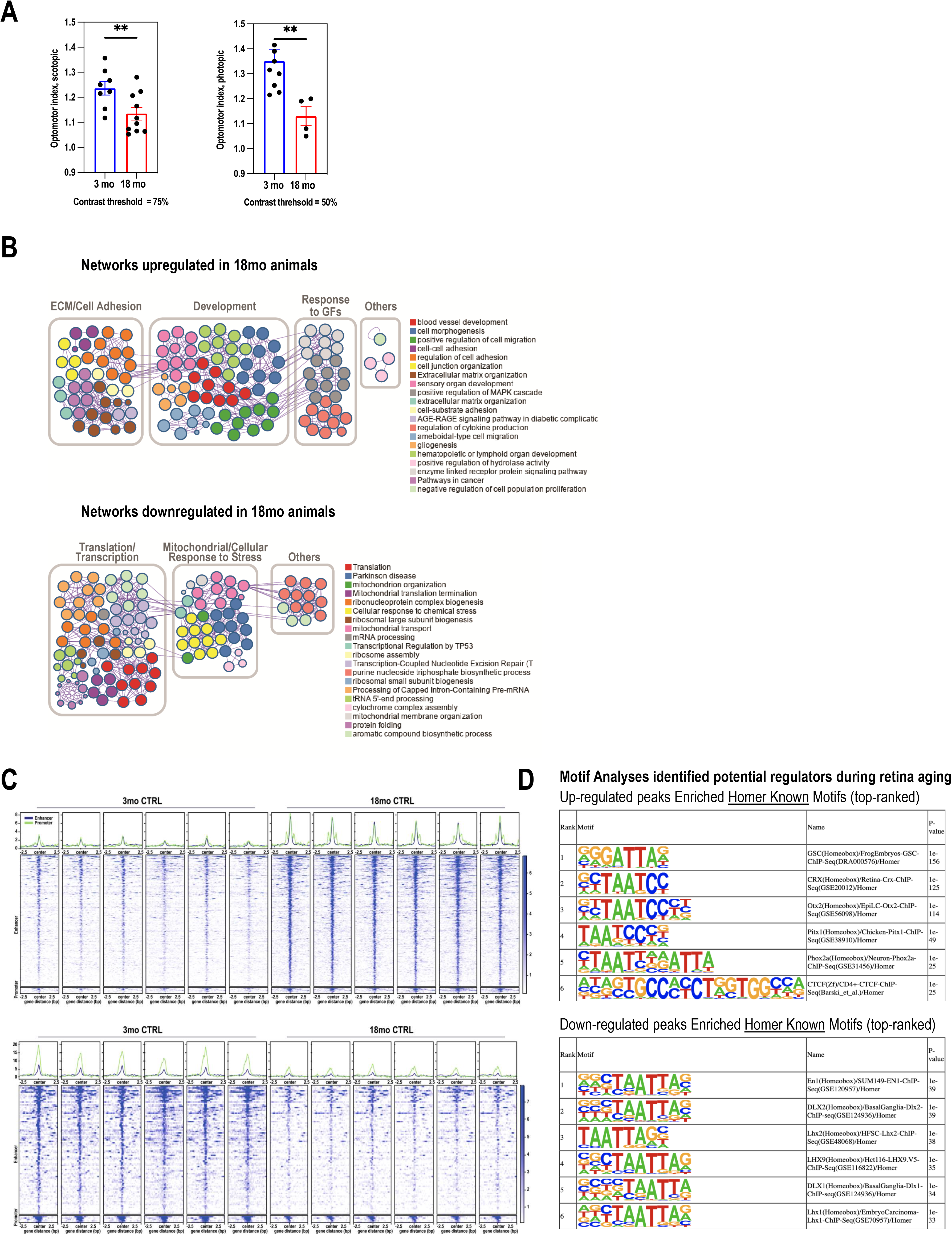
**A**. Summary of OMRs at various pattern contrasts. qOMR in scotopic conditions at 75% pattern contrast (left) 3-month-old mice (mean, 1.236 ± 0.078 [SD]; *n* = 8 animals) and 18-month-old mice (mean, 1.134 ± 0.08; *n*=10 animals) (t=2.73, p=0.0076). qOMR in photopic conditions at 50% pattern contrast. 3-month-old mice (mean, 1.35 ± 0.16 [SD]; *n* = 9 animals) and 18-month-old mice (mean, 1.13 ± 0.077; *n*=4 eyes) (t=3.58, p=0.0047), (Welch’s unpaired, two-tailed t-test). **B**. Metascape networks of enriched pathway from up- and down-regulated DEGs during aging. Each node in the figure represents one enriched ontology term. Node size indicates the number of genes in each term. Node color represent its cluster identity and nodes of the same color belong to the same cluster. One term from each cluster is selected to have its term description shown as label in the figure. All the enriched terms were further bounded and annotated as more general classes. **C**. Profile plots and heatmap plots show up- and down-regulation during aging (3-month versus 18-month) in chromatin accessibility (deepTools normalized RPKM with bin size 5) with 2.5Kb flanking distance to center of each region. n = 6 biological replicates. **D**. Motif Analyses performed by using Homer tool identified potential regulators in response to mild IOP in 18-month-old mouse retina from gained accessible regions. The differential analysis was performed by MACS2 with cutoff of q-value at 0.05 and with log2foldchange > 1. n = 6 biological replicates. Top-ranked enriched Homer Known motifs in p-value are listed in the table.

**Supplementary Figure 2.**
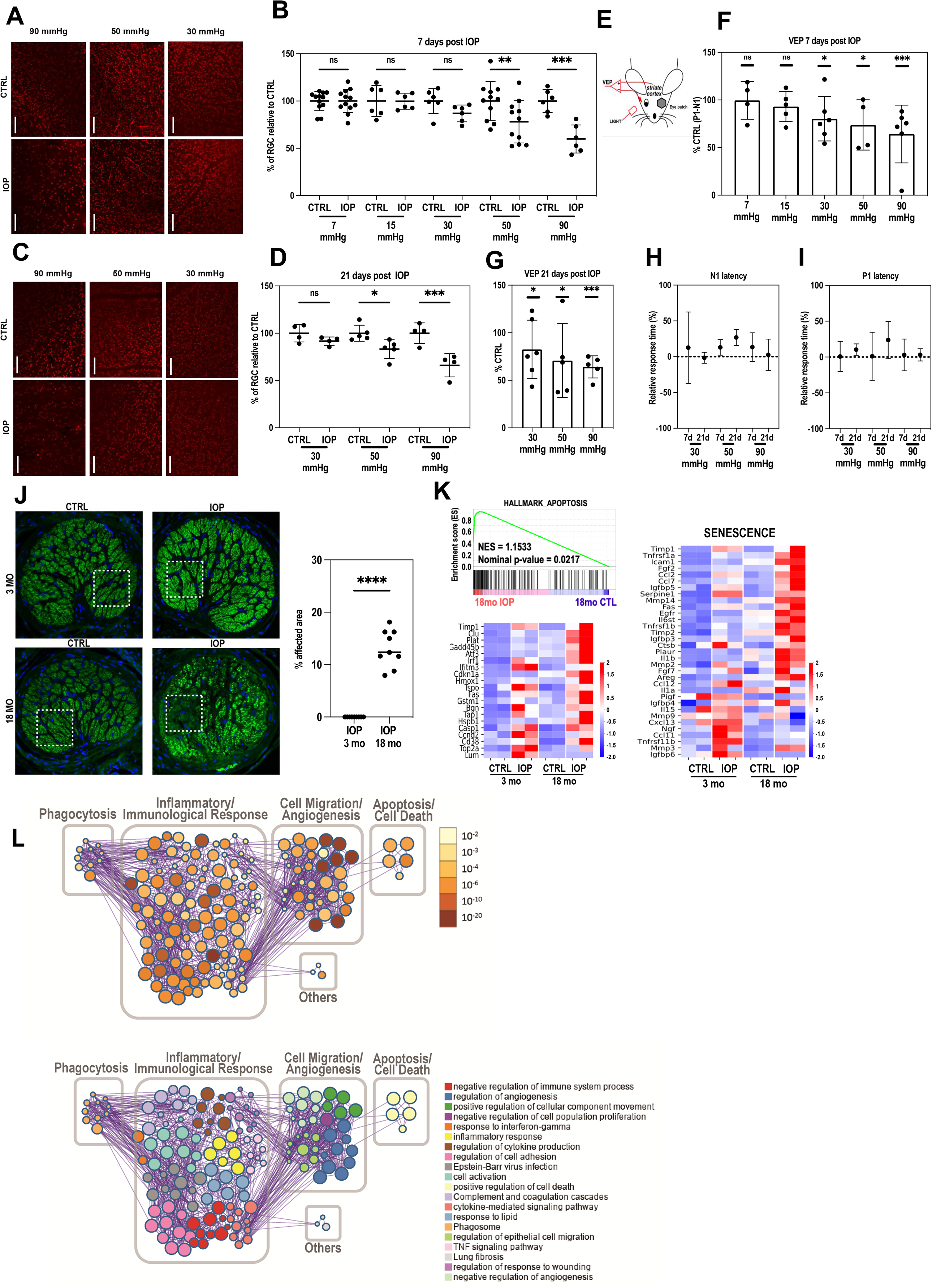
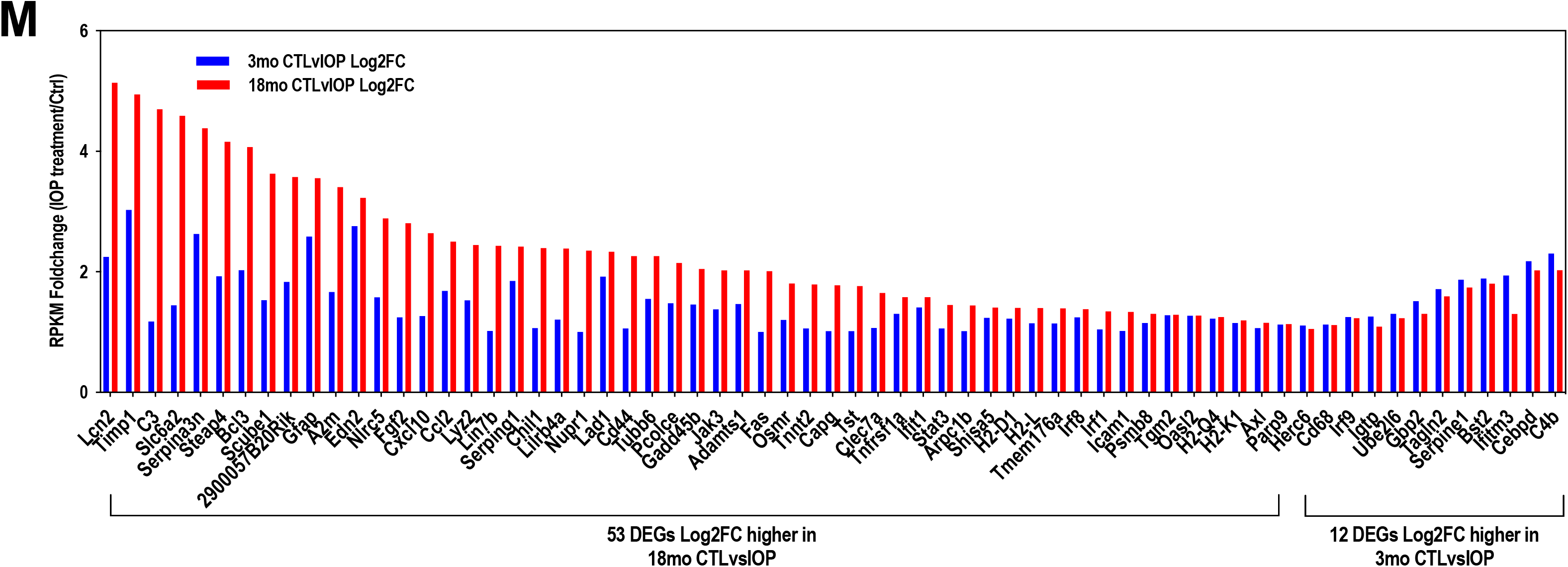
**A. and C**. Representative micrographs from Brn3a-stained retinal flat mounts, RGC count 7 days post-IOP elevation (A) and 21 days post-IOP elevation (C). Scale bar = 100 μm. **B and D**. Quantification of RGC count from retinal micrographs in IOP-treated and non-treated retinas in 3-month, old C57BL/6J mice 7 days post-IOP elevation (D) and 21 days post-IOP elevation. Eyes were exposed to 30, 50, or 90 mmHg pressure for 1h. 7 days post-IOP cohort was exposed to 7 and 15mmHg pressure for 1h. Each data point is an average of 6 independent frames in the central and peripheral retina, represented as % RGC count as compared to contralateral healthy eye (* p < 0.05, *** p < 0.005). (one-way ANOVA, multiple comparisons). Visual Evoked Potential measurements. **E**. Scheme of the VEP reading, with the measurement electrode in the striate cortex, and the reference in the mouth; measurements were taken unilaterally, with contralateral eye covered with an eye patch. **F**. Quantification of the P1-N1 visual potential from the VEP traces 7 days post-procedure in response to exposure to 7, 15, 30, and 90 mmHg. **G**. Quantification of the P1-N1 visual potential from the VEP after 21 days. **H**. Relative change in the VEP N1 latency in response to different levels of hypertension. I. Relative change in the VEP P1 latency in response to different levels of hypertension. Error bars represent SEM. **I**. *In situ* RNA hybridization staining by RNAscope in focused cell layers of retina in 18 month mice 2 days after mild IOP elevation. **J**. ONH images. Dashed line indicates area used for neurofilament count. **K**. GSEA enrichment plot of hallmark apoptosis genes and row Z-score heatmap of both hallmark apoptosis and ECM genes in 3-month and 18-month retinae upon IOP stress versus contralateral control. **L**. Metascape networks of enriched pathways from up-regulated DEGs upon IOP treatment in 18-month-old mouse retinae. The enrichment network with nodes colored by FDR q-value were shown by a discrete color scale (see legend) in the upper figure. Each node in the figure represents one enriched ontology term in the bottom. Node size indicates the number of genes in each term. Node color represent its cluster identity and nodes of the same color belong to the same cluster. One term from each cluster is selected to have its term description shown as label in the figure. All the enriched terms were further bounded and annotated as more general classes.

**Supplementary Figure 3.**
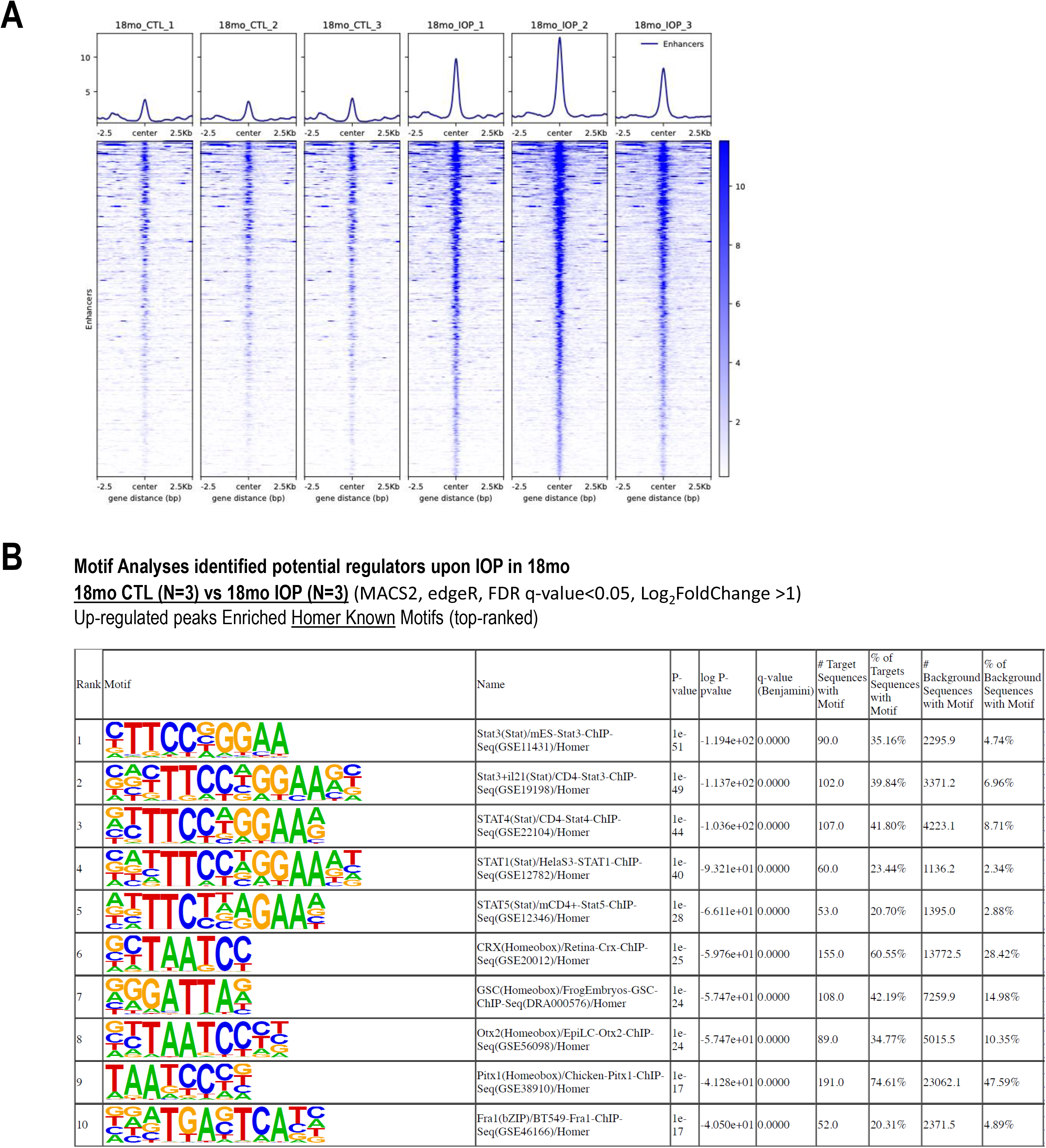
**A**. Profile plots and heatmap plots for chromatin accessibility intensities (deepTools normalized RPKM with bin size 5) over all up-regulated enhancer regions (2.5Kb flanking distance to center of each region). **B**. Motif Analyses performed by using Homer tool identified potential regulators in response to mild IOP in 18-month-old mouse retina from gained accessible regions. The differential analysis was performed by MACS2 with cutoff of q-value at 0.05 and with log2foldchange > 1. n = 3 biological replicates. Top-ranked enriched <u>Homer Known</u> motifs in p-value are listed in the table.

**Supplementary Figure 4.**
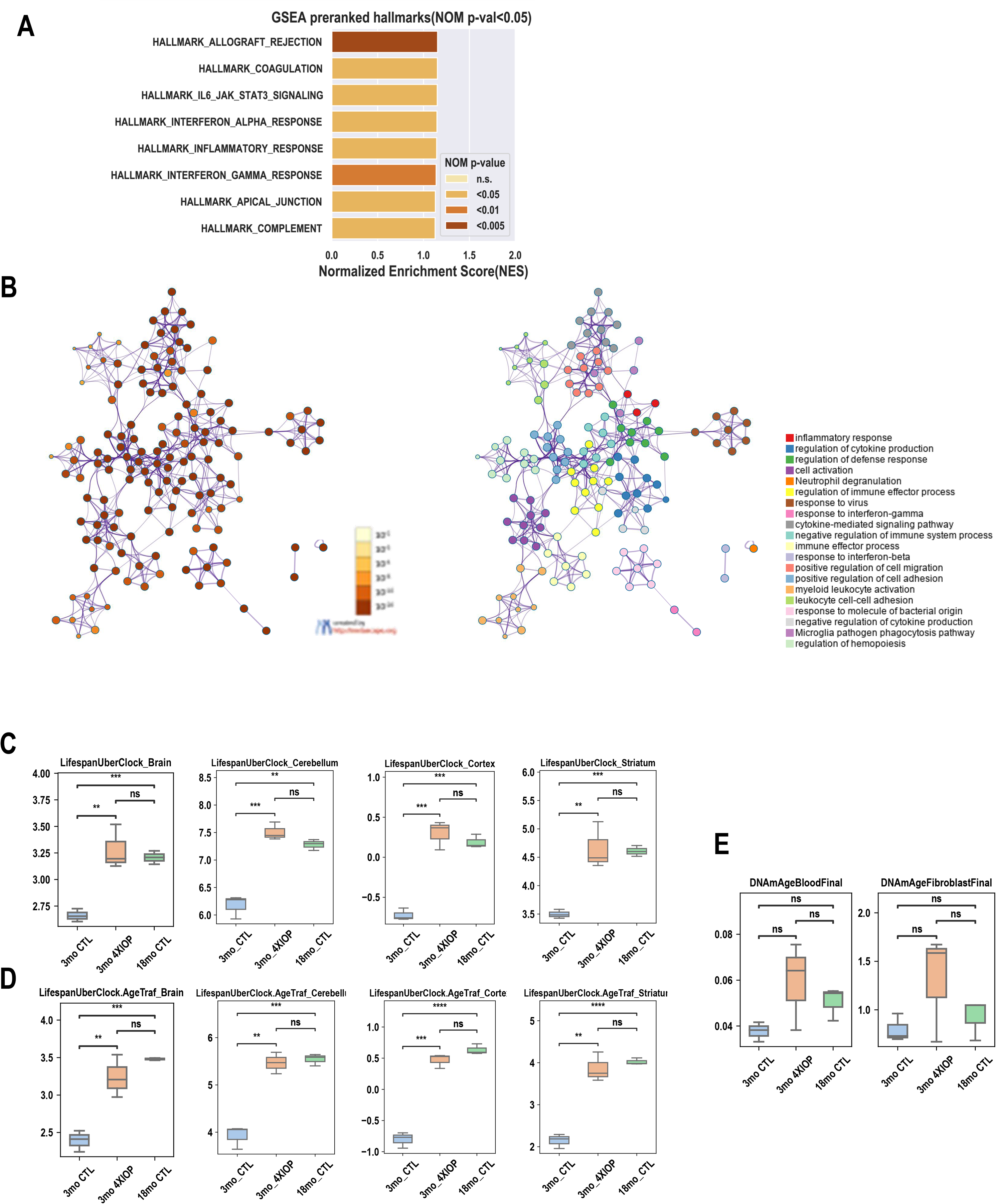
**A**. The bar plot showed normalized enrichment scores (NES) of all defined significant enriched (nominal p-value <5%) hallmark gene sets from pre-ranked GSEA in 3-month-old wildtype mouse retinae upon multiple IOP treatments. n = 2 biological replicates. NES determines the magnitude of enrichment of each hallmark gene set across all analyzed gene sets. The statistical significance in nominal (NOM) p-value were indicated by a discrete color scale. **B**. Metascape networks of enriched pathway from up-regulated DEGs in 3-month-old mouse retinae upon multiple IOP treatments. Each node in the figure represents one enriched ontology term. Node size indicates the number of genes in each term. Node color represent its cluster identity and nodes of the same color belong to the same cluster. One term from each cluster is selected to have its term description shown as label in the figure. **C**. Predicted age (by unit of years) by aging clock microarray analysis of one-time IOP treated 3-month and 18-month retinae and multiple IOP treated 3-month retinae. n = 3 biological replicates.

## Notes

### Competing Interest Statement

The authors have declared no competing interest.

### Summary of Updates

Conversion to pdf erased some symbols on figures. Current version has all symbols and annotations.

